# *De Novo* RNA Tertiary Structure Prediction at Atomic Resolution Using Geometric Potentials from Deep Learning

**DOI:** 10.1101/2022.05.15.491755

**Authors:** Robin Pearce, Gilbert S. Omenn, Yang Zhang

**Affiliations:** Department of Computational Medicine and Bioinformatics, University of Michigan, Ann Arbor, MI 48109 USA; Department of Biological Chemistry, University of Michigan, Ann Arbor, MI 48109, USA; Departments of Internal Medicine and Human Genetics and School of Public Health, University of Michigan, Ann Arbor, MI 48109, USA

## Abstract

Experimental characterization of RNA structure remains difficult, especially for non-coding RNAs that are critical to many cellular activities. We developed DeepFoldRNA to predict RNA structures from sequence alone by coupling deep self-attention neural networks with gradient-based folding simulations. The method was tested on two independent benchmark datasets from Rfam families and RNA-Puzzle experiments, where DeepFoldRNA constructed models with an average RMSD=2.69 Å and TM-score=0.743, which outperformed state-of-the-art methods and the best models submitted from the RNA-Puzzles community by a large margin. On average, DeepFoldRNA required ~1 minute to fold medium-sized RNAs, which was ~350-4000 times faster than the leading Monte Carlo simulation approaches. These results demonstrate the major advantage of advanced deep learning techniques to learn more accurate information from evolutionary profiles than knowledge-based potentials derived from simple statistics of the PDB library. The high speed and accuracy of the developed method should enable large-scale atomic-level RNA structure modeling applications.

## INTRODUCTION

RNAs are vital macromolecules that play a fundamental role in many cellular processes in living organisms, including mediating gene translation, serving as catalysts of important biological reactions, and regulating gene expression^1^. Many of these functions are determined by their unique three-dimensional structures, which in turn are dictated by their nucleic acid sequences. Although understanding RNA structures is fundamental to elucidating their functions, there is an enormous discrepancy between the number of known RNA sequences and the number of solved structures. For example, while >31 million RNA sequences have been deposited in the RNAcentral database^2^, there are <500 non-redundant RNA structures solved in the Protein Data Bank (PDB) at a resolution of ~2 Å and <30 are composed of >70 nucleotides. Furthermore, only 99 of the 4,192 Rfam families have members with solved structures^3^. Thus, there is an urgent need to develop computational RNA structure prediction methods capable of addressing this stark disparity.

The goal of RNA structure prediction is to determine the spatial location of every atom in an RNA molecule starting from its nucleic acid sequence. Some state-of-the-art methods take a physics-based approach to model RNA structures by identifying low free-energy states through Monte Carlo simulations^4^, while others approach the problem by assembling homologous fragments for a given nucleic acid sequence guided by knowledge-based energy functions^5^. However, even with the assistance of human expert intervention and experimental data, these methods struggle to produce accurate folds for larger, more complex RNA molecules, rarely achieving RMSDs lower than 8-12 Å in blind RNA structure prediction studies^6,7^. Moreover, the results are typically worse for automatic modeling methods, which may produce models with around 20 Å RMSDs for complex folds^4^. Progress has been made by using deep learning to predict secondary structure and contact information to guide the folding simulations^8–11^; however, the improvements remain unsatisfactory and current state-of-the-art methods rarely achieve atomic resolution models, i.e., <2Å RMSD^12^, for complex RNA folds^5–7,13^. Recently, deep learning approaches have been successfully applied to the problem of model selection^14^. Nevertheless, the success of these methods is predicated on generating conformations that are close to the native structures, where atomic resolution was only obtained after utilizing restraints from native structures, which are not available in practical modeling applications.

To improve the performance of RNA structure prediction methods, we drew inspiration from recent advances in protein structure prediction, where deep learning techniques have revolutionized the field^15–17^. Toward this goal, we developed DeepFoldRNA, which uses a self-attention-based neural network architecture to predict geometric restraints, where 3D RNA structures are then built using limited-memory Broyden-Fletcher-Goldfarb-Shanno (L-BFGS) minimization simulations. Across multiple test experiments, DeepFoldRNA drastically outperformed other state-of-the-art modeling methods and consistently achieved atomic-level resolution for complex RNA folds. In addition, due to the rapid gradient-based folding simulations, RNAs could be folded in a tiny fraction of the time required by current methods. The speed and accuracy of DeepFoldRNA will allow for large-scale elucidation of RNA structure and function, addressing a fundamental problem in structural biology. Each component of the program, including the deep learning models and L-BFGS optimization pipeline, are integrated into a stand-alone package at https://github.com/robpearc/DeepFoldRNA and an online webserver is available at https://zhanggroup.org/DeepFoldRNA, from which users can generate structure models for their own RNA of interest.

## RESULTS AND DISCUSSION

DeepFoldRNA is a method for fully-automated RNA structure prediction that consists of two consecutive modules (Figure 1). In the restraint generation module (Figure 1A), multiple sequence alignments (MSAs) of RNAs are collected by iteratively searching through multiple nucleic acid sequence databases using rMSA^18^, where spatial restraints, including pairwise distance and inter-residue/backbone torsion angles maps, are predicted using self-attention neural networks that are built on two transformer elements with information encoded from the sequence, MSA, and pairwise positional embeddings. In the structure construction module (Figure 1B), the predicted geometric restraints are converted into composite potentials by taking the negative log-likelihood of the binned probability predictions, which are then used to guide the L-BFGS folding simulations.

**Figure 1.**
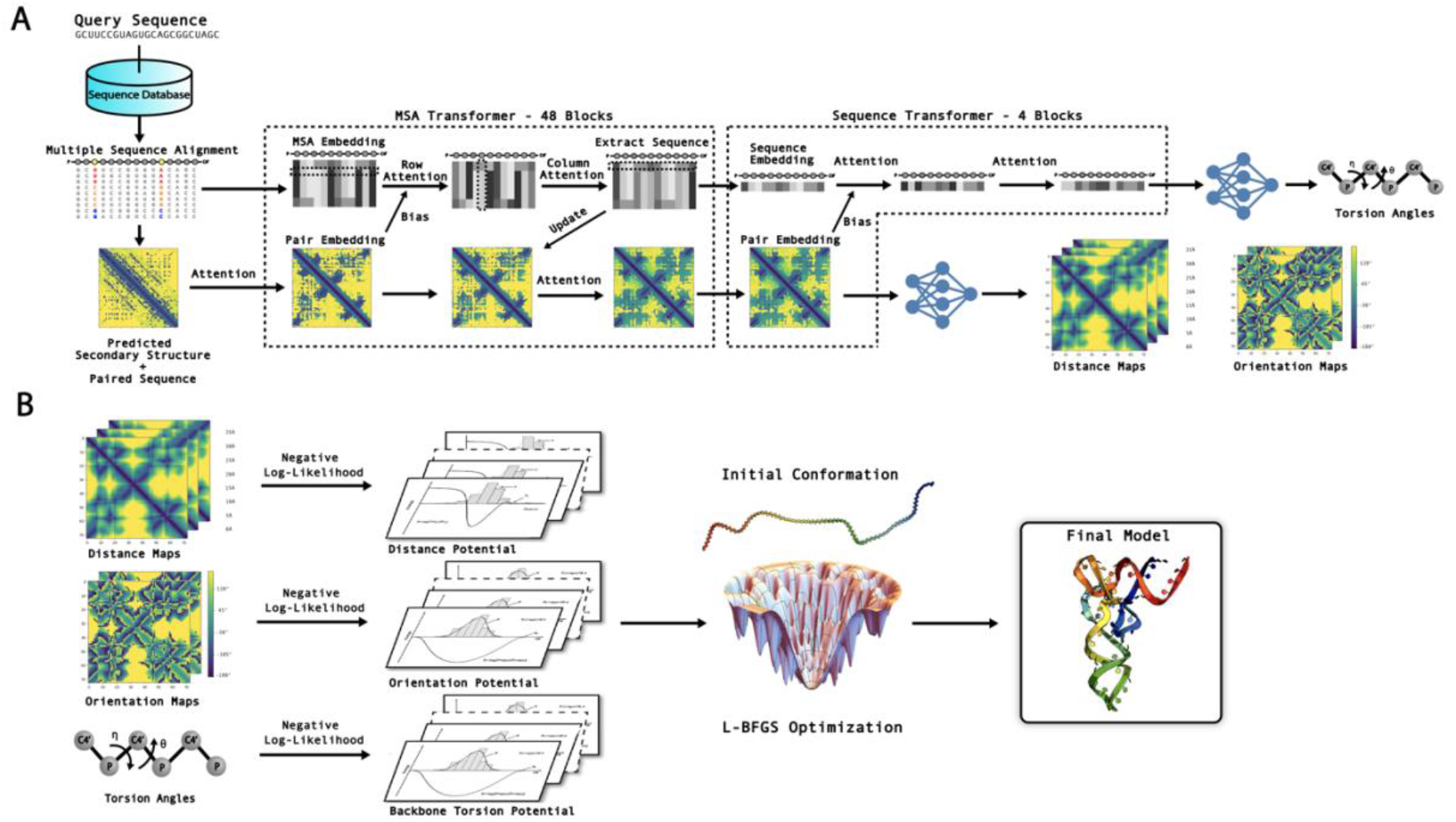
Overview of the DeepFoldRNA pipeline. A) Starting from a nucleic acid sequence, multiple RNA sequence databases are searched to create a multiple sequence alignment (MSA) for the query RNA, which is embedded into the network to initialize the MSA representation. The raw MSA is also used to derive the secondary structure prediction and initialize the pair embedding. The MSA and pair embeddings are then processed by the MSA Transformer layers, which use multiple self-attention mechanisms to refine the initial embeddings, where communication is encouraged between the two to ensure consistency. Next, the sequence embedding is extracted from the row in the final MSA embedding corresponding to the query sequence, which is further processed using self-attention mechanisms by the Sequence Transformer layers. Finally, the distance and inter-residue torsion angle maps are predicted from a linear projection of the final pair embedding, while the backbone pseudo-torsion angles are generated by a linear projection of the sequence embedding. B) The geometric restraints are converted into a negative-log likelihood potential to guide the L-BFGS simulations for final RNA model construction.

Two datasets were constructed to test DeepFoldRNA. The first was collected from Rfam families^3^ with experimentally solved structures, where we curated a set of 4082 Rfam structures with complex folds and lengths between 70-250 nucleotides. From this set, we obtained 105 non-redundant RNA structures from 32 Rfam families after using a sequence identity cutoff of 80%. The second dataset was taken from the community-wide RNA-Puzzles experiment^6,7,13,19^ and consisted of 17 non-redundant, monomeric RNA structures where the models predicted by all groups were available to be downloaded at https://github.com/RNA-Puzzles/standardized_dataset. All targets in the test sets, together with those at >80% sequence identity to them, were held out from training the DeepFoldRNA pipeline.

### DeepFoldRNA accurately predicts geometric restraints

Two central geometric restraints are predicted by DeepFoldRNA, including distance and orientation maps. The distance maps include the pairwise distances between the nitrogen atoms of the base bonded to the ribose sugar (N1 for pyrimidines and N9 for purines) as well as the backbone C4’ and P atoms (Figure S1A), while the inter-residue orientations include 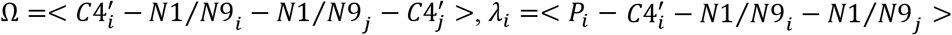, and 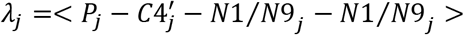, where *i* and *j* are the nucleotide indices along the sequence (Figure S1B). The network of Module-1 generates probability distributions for each of the geometric restraints, where the distances and orientations are divided into 40 and 25 bins, respectively (see Methods).

To assess the accuracy of the predicted restraints, we list in Table 1 the Mean Absolute Errors (MAEs) for the top *L*, 5*L* and 10*L* medium/long-range (|*i* – *j*|>12) distance and orientation restraints predicted by DeepFoldRNA for the 122 RNAs in the two test sets. Here, 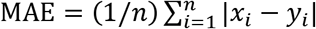, where *x_i_* is the value of the predicted restraint with the maximum probability score for a selected residue pair, *y_i_* is the corresponding value in the native structure, and *n* is the number of restraints considered. Since to our knowledge there is no comparable distance/orientation prediction method for RNA, as a control we also list the distance/orientation parameters taken from the predicted models by two state-of-the-art modeling methods: SimRNA^4^ and Rosetta FARFAR2^5^, which have been among the most accurate automatic modeling approaches in previous RNA-Puzzles experiments^6,7,13^. To provide a fair comparison between the methods, the predicted secondary structures used by DeepFoldRNA were used as constraints during the SimRNA and FARFAR2 simulations, where the exact procedures used to run both programs are provided in Supplementary Texts S1 and S2. Overall, DeepFoldRNA produced accurate distance and orientation predictions, where the average MAEs for the top *L*, 5*L* and 10*L* N1/N9 distances were 0.72, 0.83 and 0.93 Å, respectively, which were ~9-11 times lower than those extracted from the SimRNA and FARFAR2 models. For the *Ω/λ* orientations, the average *L*, 5*L* and 10*L* MAEs were 0.17/0.14, 0.20/0.16 and 0.23/0.17 radians, respectively, which were around 4-6.5 times lower than those obtained from the SimRNA and FARFAR2 models. These data demonstrate the ability of DeepFoldRNA to create very accurate restraint predictions, which are crucial to its modeling performance.

**Table 1:**
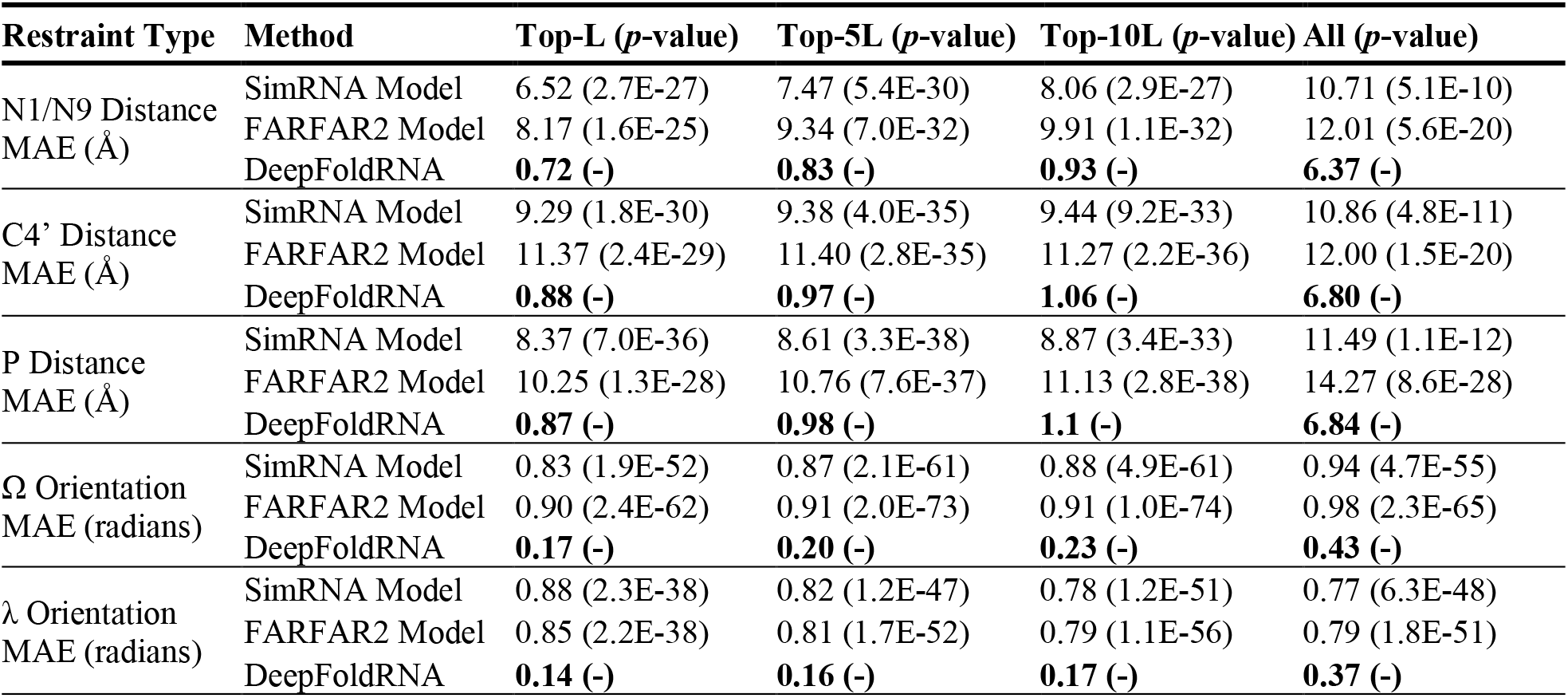
Summary of the accuracy of the DeepFoldRNA predicted restraints in terms of the Mean Absolute Errors (MAEs) for the top medium/long-range (|*i* – *j*|>12) restraints, where *L* is the RNA length. The *p*-values were calculated between DeepFoldRNA and the control methods using paired, two-sided Student’s t-tests.

| Restraint Type | Method | Top-L ($p$-value) | Top-5L ($p$-value) | Top-10L ($p$-value) | All ($p$-value) |
| --- | --- | --- | --- | --- | --- |
| N1/N9 Distance<br>MAE (Å) | SimRNA Model | 6.52 (2.7E-27) | 7.47 (5.4E-30) | 8.06 (2.9E-27) | 10.71 (5.1E-10) |
|  | FARFAR2 Model | 8.17 (1.6E-25) | 9.34 (7.0E-32) | 9.91 (1.1E-32) | 12.01 (5.6E-20) |
|  | DeepFoldRNA | **0.72 (-)** | **0.83 (-)** | **0.93 (-)** | **6.37 (-)** |
| C4’ Distance<br>MAE (Å) | SimRNA Model | 9.29 (1.8E-30) | 9.38 (4.0E-35) | 9.44 (9.2E-33) | 10.86 (4.8E-11) |
|  | FARFAR2 Model | 11.37 (2.4E-29) | 11.40 (2.8E-35) | 11.27 (2.2E-36) | 12.00 (1.5E-20) |
|  | DeepFoldRNA | **0.88 (-)** | **0.97 (-)** | **1.06 (-)** | **6.80 (-)** |
| P Distance<br>MAE (Å) | SimRNA Model | 8.37 (7.0E-36) | 8.61 (3.3E-38) | 8.87 (3.4E-33) | 11.49 (1.1E-12) |
|  | FARFAR2 Model | 10.25 (1.3E-28) | 10.76 (7.6E-37) | 11.13 (2.8E-38) | 14.27 (8.6E-28) |
|  | DeepFoldRNA | **0.87 (-)** | **0.98 (-)** | **1.1 (-)** | **6.84 (-)** |
| $\Omega$ Orientation<br>MAE (radians) | SimRNA Model | 0.83 (1.9E-52) | 0.87 (2.1E-61) | 0.88 (4.9E-61) | 0.94 (4.7E-55) |
|  | FARFAR2 Model | 0.90 (2.4E-62) | 0.91 (2.0E-73) | 0.91 (1.0E-74) | 0.98 (2.3E-65) |
|  | DeepFoldRNA | **0.17 (-)** | **0.20 (-)** | **0.23 (-)** | **0.43 (-)** |
| $\lambda$ Orientation<br>MAE (radians) | SimRNA Model | 0.88 (2.3E-38) | 0.82 (1.2E-47) | 0.78 (1.2E-51) | 0.77 (6.3E-48) |
|  | FARFAR2 Model | 0.85 (2.2E-38) | 0.81 (1.7E-52) | 0.79 (1.1E-56) | 0.79 (1.8E-51) |
|  | DeepFoldRNA | **0.14 (-)** | **0.16 (-)** | **0.17 (-)** | **0.37 (-)** |

It is noted that since the SimRNA and FARFAR2 models do not have confidence scores associated with each distance/orientation, we selected restraints based on the DeepFoldRNA confidence scores alone in the above comparison. To remove the bias in restraint selection, we present a comparison for all medium/long-range restraints in the last column of Table 1. As expected, the MAEs were much larger for the DeepFoldRNA restraints when all residues were considered, suggesting the sensitivity of the DeepFoldRNA confidence scores and the sensibility of 3D model construction based on a limited number of high-ranking restraints. Nevertheless, the MAEs for DeepFoldRNA were still considerably lower than those from the SimRNA and FARFAR2 models. Interestingly, the MAEs for the SimRNA and FARFAR2 models were also typically smaller (except for the *λ* orientation) for the top-ranking residues selected by their DeepFoldRNA confidence scores than for all residues; this is in part due to the fact that DeepFoldRNA typically has higher confidence scores in conserved regions for which SimRNA and FARFAR2 also tend to generate slightly better models.

### DeepFoldRNA dramatically outperforms state-of-the-art methods on the Rfam dataset

To evaluate the modeling performance of DeepFoldRNA, Table 2 presents a summary of the 3D modeling results on the 105 RNAs from the Rfam dataset in terms of the average/median RMSDs and TM-scores relative to the experimental structures along with the results by SimRNA and FARFAR2. Here, TM-score is a length-independent metric for assessing structural similarity that takes a value in the range [0, 1], where a TM-score=1 corresponds to an identical structural match and a TM-score >0.45 indicates that two RNAs share the same global fold^20,21^.

**Table 2:** Summary of the structure modeling results by DeepFoldRNA compared to the control methods on the 105 test RNAs from the Rfam dataset. The RMSDs and TM-scores were calculated using the RNA-align program^37^ based on sequence-dependent superposition of the C3’ atoms. The *p*-values were calculated between DeepFoldRNA and the control methods using paired, two-sided Student’s t-tests.

| Method | RMSD Avg/Median<br>($p$-value) | TM-score Avg/Median<br>($p$-value) | Correct<br>Folds* | RMSD<sub>DeepFoldRNA</sub><br><RMSD<sub>Method</sub><sup>‡</sup> |
| --- | --- | --- | --- | --- |
| SimRNA | 19.37/17.23 (6.2E-44) | 0.228/0.222 (1.0E-66) | 0.0% | 100.0% |
| FARFAR2 | 21.07/19.17 (8.4E-52) | 0.228/0.219 (6.6E-64) | 0.0% | 100.0% |
| DeepFoldRNA | **2.68/2.11 (-)** | **0.757/0.779 (-)** | **99.0%** | **-** |

\* This column represents the percent of RNAs with TM-scores $\geq 0.45$ .
‡ This column indicates the percent of test RNAs for which DeepFoldRNA generated a model with a lower RMSD than the control method.

On average, DeepFoldRNA achieved a TM-score of 0.757, which was 232% higher than that attained by SimRNA and FARFAR2 (0.228); the differences were highly statistically significant with p-values of 1.0E-66 and 6.6E-64 for the comparison with SimRNA and FARFAR2, respectively. Meanwhile, the average RMSD of the DeepFoldRNA models was 2.68 Å compared to 19.37 Å and 21.07 Å for SimRNA and FARFAR2, respectively; the differences were again highly statistically significant with p-values of 6.2E-44 and 8.4E-52. When considering the median values, DeepFoldRNA produced models with a median RMSD of 2.11 Å (SimRNA: 17.23 Å; FARFAR2: 19.17 Å) and a median TM-score of 0.779 (SimRNA: 0.222; FARFAR2: 0.219), corresponding to close atomic matches between the predicted and native structures.

In Figures 2A-B and D-E, we present head-to-head RMSD and TM-score comparisons of DeepFoldRNA with SimRNA and FARFAR2. Overall, DeepFoldRNA generated models with lower RMSDs and higher TM-scores than the control methods for all of the test RNAs. Furthermore, Figures 2C and 2F list the number of models produced below a specific RMSD or above a given TM-score threshold. When considering a cutoff TM-score of 0.45, for example, DeepFoldRNA generated correct global folds for 99% or all but one of the test RNAs, while the control methods were unable to generate correct global folds for any of the targets. DeepFoldRNA also consistently generated models with atomic-level accuracy, where 46 of the 105 models (43.8%) had RMSDs <2 Å to their experimental structures. When considering a more permissive RMSD cutoff of <4.0 Å to define a native-like structure, 86.7% of the DeepFoldRNA models met this criterion, while none of the models by the control methods did so.

**Figure 2.**
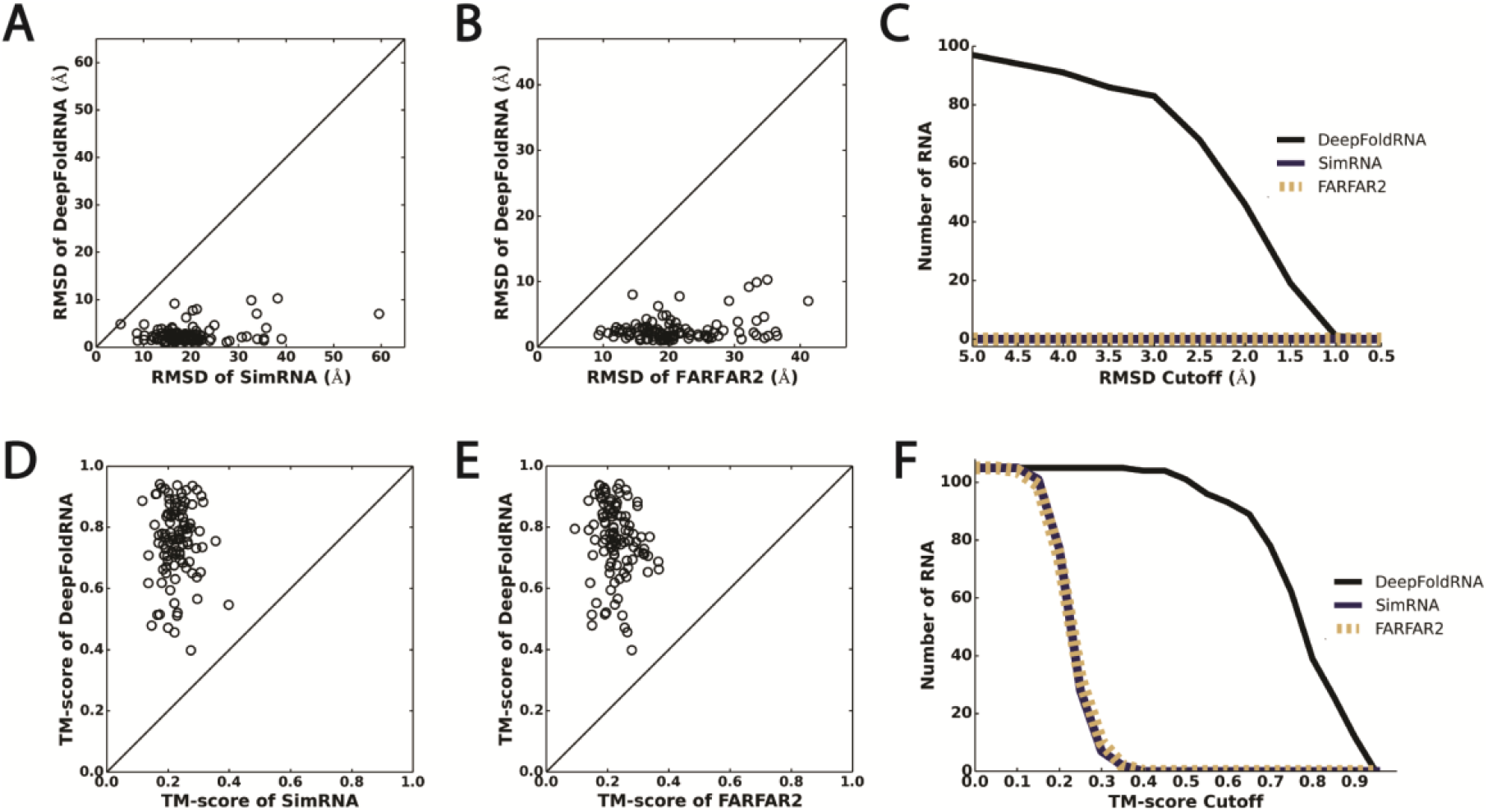
Head-to-head RMSD and TM-score comparisons of DeepFoldRNA with the selected state-of-the-art methods on the 105 Rfam RNA strucures. A) RMSD comparison with SimRNA, B) RMSD comparison with FARFAR2, C) Number of targets below a given RMSD threshold, D) TM-score comparison with SimRNA, E) TM-score comparison with FARFAR2, F) Number of targets above a given TM-score threshold.

Importantly, the success of DeepFoldRNA modeling was not limited to any specific fold type. Figure 3 plots representative models across all 32 Rfam families in the test set. For 14 of the 32 families (43.8%), DeepFoldRNA generated atomic resolution models with <2Å RMSD and 100% of the models possessed correct global folds with TM-scores >0.45. Highly accurate models could be constructed for well represented families such as RF00001 (composed of 5S ribosomal RNAs) and RF00005 (made up of tRNAs), where the DeepFoldRNA models had RMSDs of 1.08Å and 1.09Å, respectively, corresponding to very close atomic matches between the predicted and experimental structures. Accurate models were also constructed for families with few sequence homologs. For instance, RF01689 (PDB ID 4frg, chain B, residues 1-83) is composed of AdoCbl variant RNAs and the generated MSA had relatively few homologous sequences with a Neff value (number of effective sequences) of 3.65, where DeepFoldRNA created an accurate model for this family with an RMSD of 2.12 Å and a TM-score of 0.709.

**Figure 3.**
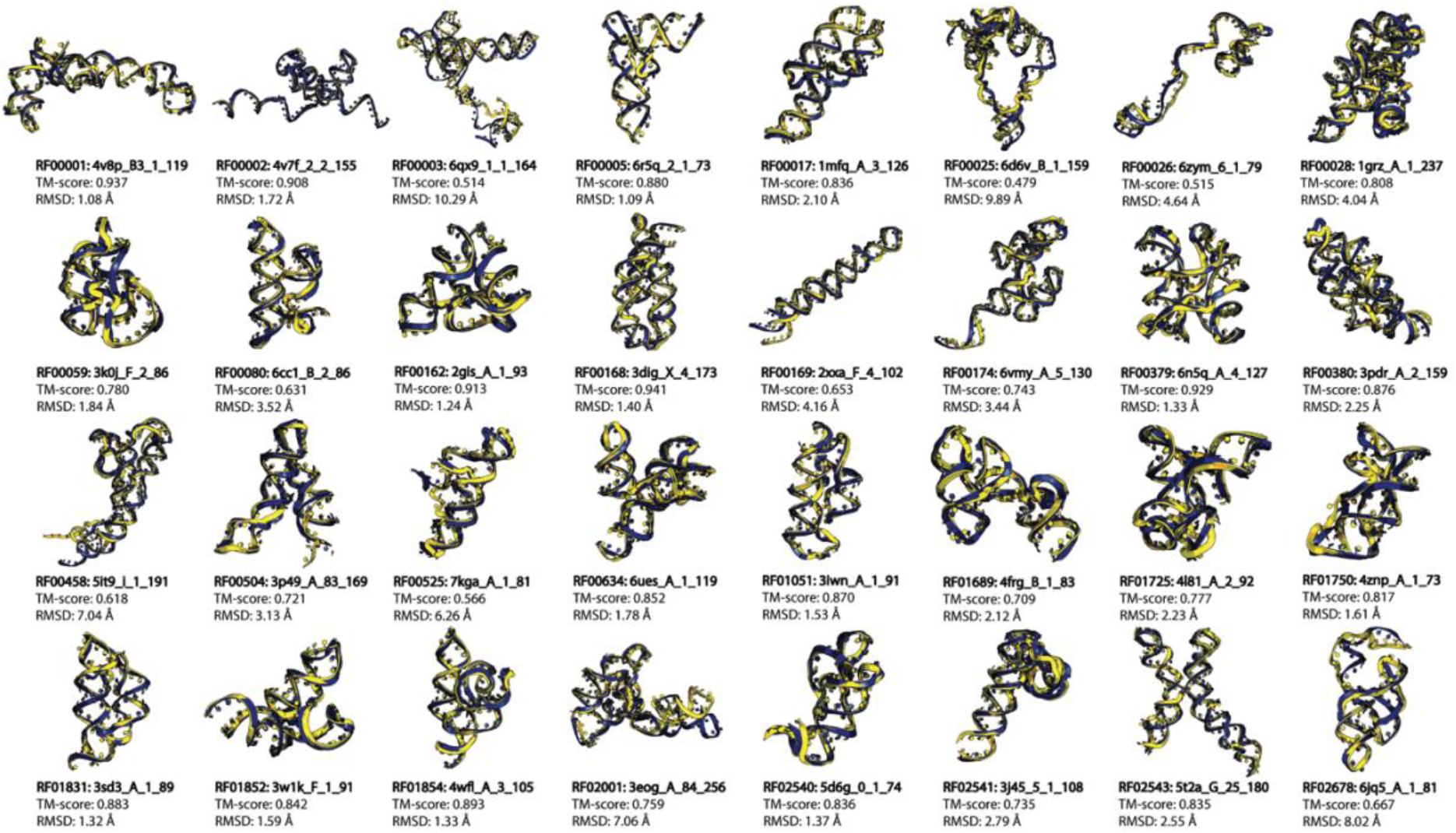
Representative models generated by DeepFoldRNA for each of the 32 Rfam families. The modeled structures in blue are superposed with the native structures in yellow. The PDB IDs, chain ids, and residue numbers are shown below each RNA together with the TM-scores and RMSDs.

Interestingly, for models with higher RMSDs, the modeling errors were often localized in flexible or unstructured regions of the RNAs. For instance, for RF02678 (PDB ID 6jq5, chain A, residues 1-81) the model generated by DeepFoldRNA had an RMSD of 8.02Å, where the deviation between the modeled and native structures was mainly confined to the unpaired region of the structure from residues 64-81 (Figure S2). In the core region of the RNA (residues 1-63), in contrast, the RMSD between the modeled and native structures was only 1.40 Å, resulting in a correct global fold with a TM-score of 0.667. Overall, the results demonstrate that DeepFoldRNA is able to consistently generate correct global folds, frequently with atomic-level resolution, for RNAs across various complex fold types, drastically outperforming the leading Monte Carlo simulation methods.

### DeepFoldRNA outperforms the best models from the RNA-Puzzles community by a large margin

To further examine DeepFoldRNA with the state of the art, we tested it on 17 challenging RNA targets from the community-wide RNA-Puzzles experiment, where many of the targets lacked structural and sequence homologs^6,7,13,19^. The experiment is split into Human and Server Sections, where each group is allowed to submit up to 10 models for each target. Traditionally, the automated Server methods, which are given 48 hours to model a target, have been unable to achieve the same performance as Human groups, who are typically given 3-6 weeks and often utilize extensive expert intervention during the modeling process and restraints from fast-track experimental data^6,7,13^. Figure 4 summarizes the modeling results of DeepFoldRNA compared to all RNA-Puzzles participants.

**Figure 4.**
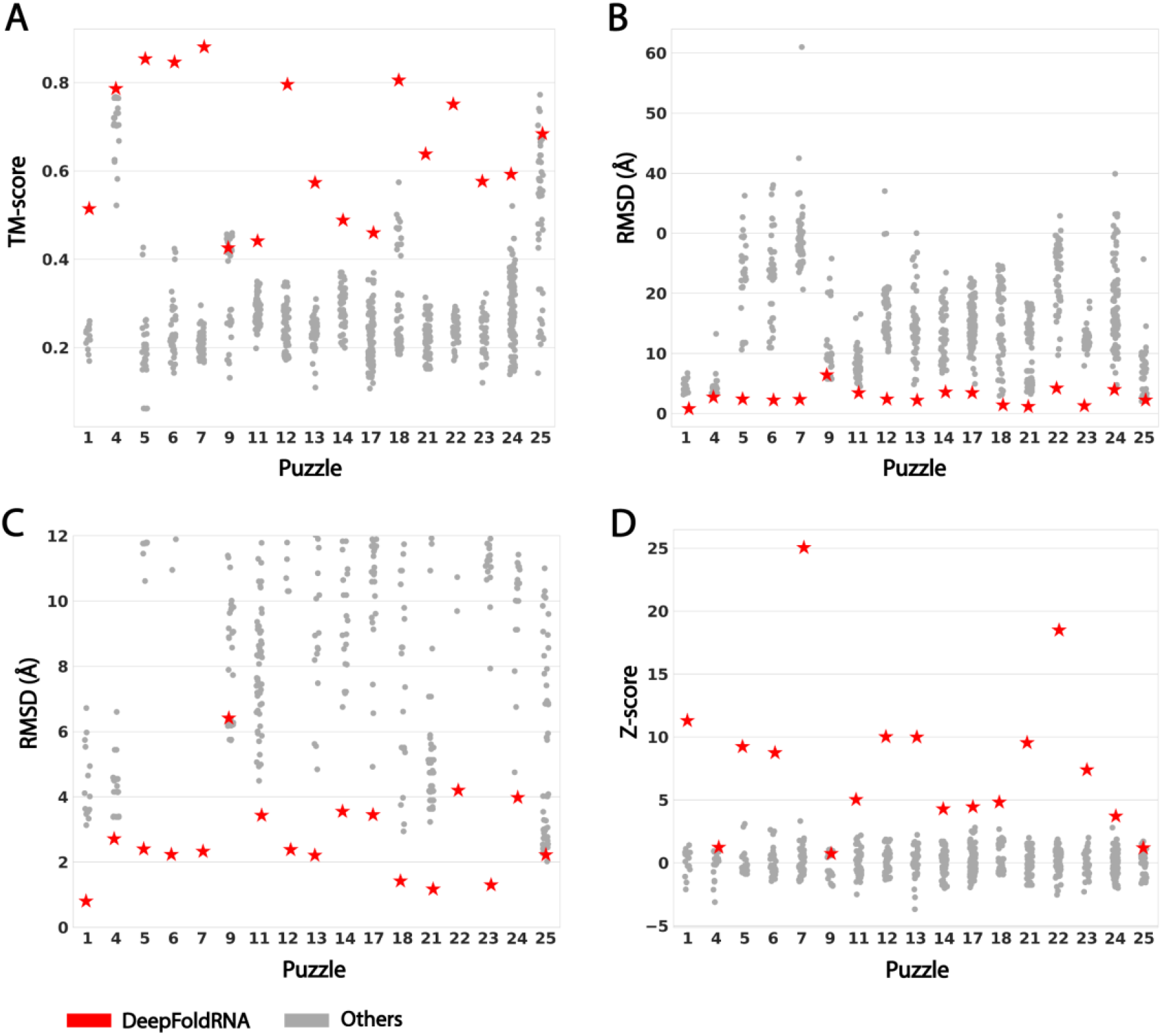
DeepFoldRNA modeling results on the 17 RNA-Puzzles targets compared to the participants. (A) TM-score; (B) RMSD; (C) Same as (B) but only for models with RMSDs below 12 Å. (D) Z-score of the TM-score for DeepFoldRNA compared to the participating groups. The RMSDs and TM-scores were calculated using the RNA-align program^37^ based on sequence-dependent superposition of the C3’ atoms.

Overall, DeepFoldRNA achieved an average TM-score of 0.654, which was 77.7% higher than the average TM-score of the first models generated by the best-performing group in RNA-Puzzles (Das Group, TM-score=0.368). If we select the best model submitted for each target by all of the RNA-Puzzles participants, the DeepFoldRNA TM-score was still 55.0% higher than that of the best models by the community (TM-score=0.422). Similarly, the average RMSD of DeepFoldRNA (2.72 Å) was 4.18 Å lower than the average RMSD of the best models (6.90 Å) generated by all RNA-Puzzles groups. When considering a cutoff TM-score of 0.45, DeepFoldRNA generated correct global folds for 15 of the 17 Puzzles (88.2%), while correct global folds could only be constructed for 5 of the 17 targets (29.4%) by the RNA-Puzzles community. Meanwhile, DeepFoldRNA generated models with <2.5 Å RMSD for 10 of the 17 cases (58.8%), while this accuracy was achieved for only one target (Puzzle 25) by the community.

Since many of the RNA-Puzzles targets lack sequence homologs, it is of interest to examine the modeling performance in relation to the quality of the generated MSAs. In Figure S3A, we plot the TM-score of the DeepFoldRNA models against the logarithm of the MSA Neff value on the RNA-Puzzles dataset. From the figure, it can be seen that there is essentially no correlation (ρ=-0.001) between the model TM-score and the MSA Neff value, suggesting that DeepFoldRNA is a robust method for the hardest class of RNA targets, which lack homologous sequence information. Furthermore, Figure S3B plots the model TM-scores vs. the MSA Neff values across the targets in both the RNA-Puzzles and Rfam datasets. Again, only a very weak correlation (ρ=0.013) existed between the Neff value and the model quality by DeepFoldRNA. Overall, these results demonstrate that DeepFoldRNA is capable of accurately folding very challenging modeling targets using a fully automated pipeline, significantly outperforming approaches from the RNA-Puzzles challenge, where many of the predictions were guided by human expert intervention and experimental restraints.

### Case studies reveal DeepFoldRNA’s ability to fold challenging targets with complex structures

A closer examination of Figure 4 shows that DeepFoldRNA achieved the best models with the highest TM-scores and lowest RMSDs for 15 out of the 17 RNA targets. If we define a Z – score = (TM_D_ – 〈TM〉)/*σ*, where TM_D_ is the TM-score of the DeepFoldRNA model, 〈TM〉 is the average TM-score of all groups and *σ* is the standard deviation, DeepFoldRNA generated a model that was better than any other submitted model by a large margin for 10 cases (i.e., Puzzles 1, 5, 6, 7, 11, 12, 13, 21, 22, and 23) with Z-scores above 5 (Figure 4D). There were only two targets (PZ9 and PZ25) for which the DeepFoldRNA model was marginally worse with RMSDs that were 0.2 and 0.66 Å higher than the best models from the RNA-Puzzles community, respectively.

In Figure 5, we present four case studies for which DeepFoldRNA achieved near-native quality models with RMSDs <2.5 Å, while all models submitted by the RNA-Puzzles community had RMSDs above 10 Å. First, **Puzzle 5** is a 188-nucleotide long lariat-capping ribozyme (PDB ID: 4p9r) that catalyzes reactions involving the formation of a 3 nucleotide 2’,5’ lariat^22^. The RNA possesses a unique open ring structure formed by the interaction between the two peripheral helical regions. The highest TM-score model submitted by the RNA-Puzzles community had a TM-score of 0.426 and an RMSD of 10.61Å, where the open ring structure was not reproduced by any of the submitted models^13^. For this target, the generated MSA by rMSA contained 17 sequence homologs, where only 3 sequences were aligned to the query with a coverage >50%, resulting in a low Neff value of 0.65. Nevertheless, the deep learning module generated accurate spatial restraints with MAEs for the top 5*L* N-N distances and *Ω/λ* orientations of 0.99 Å and 0.17/0.14 radians, respectively. Additionally, the structure produced by the folding simulations closely converged to the predicted restraints with an MAE of 0.72 Å between the predicted top 5*L* N-N distances and the model distances. This resulted in a high-quality 3D structure with a TM-score of 0.851 and RMSD of 2.43 Å, accurately recapitulating the open ring structure and again highlighting the ability of DeepFoldRNA to model challenging targets with few homologous sequences.

**Figure 5.**
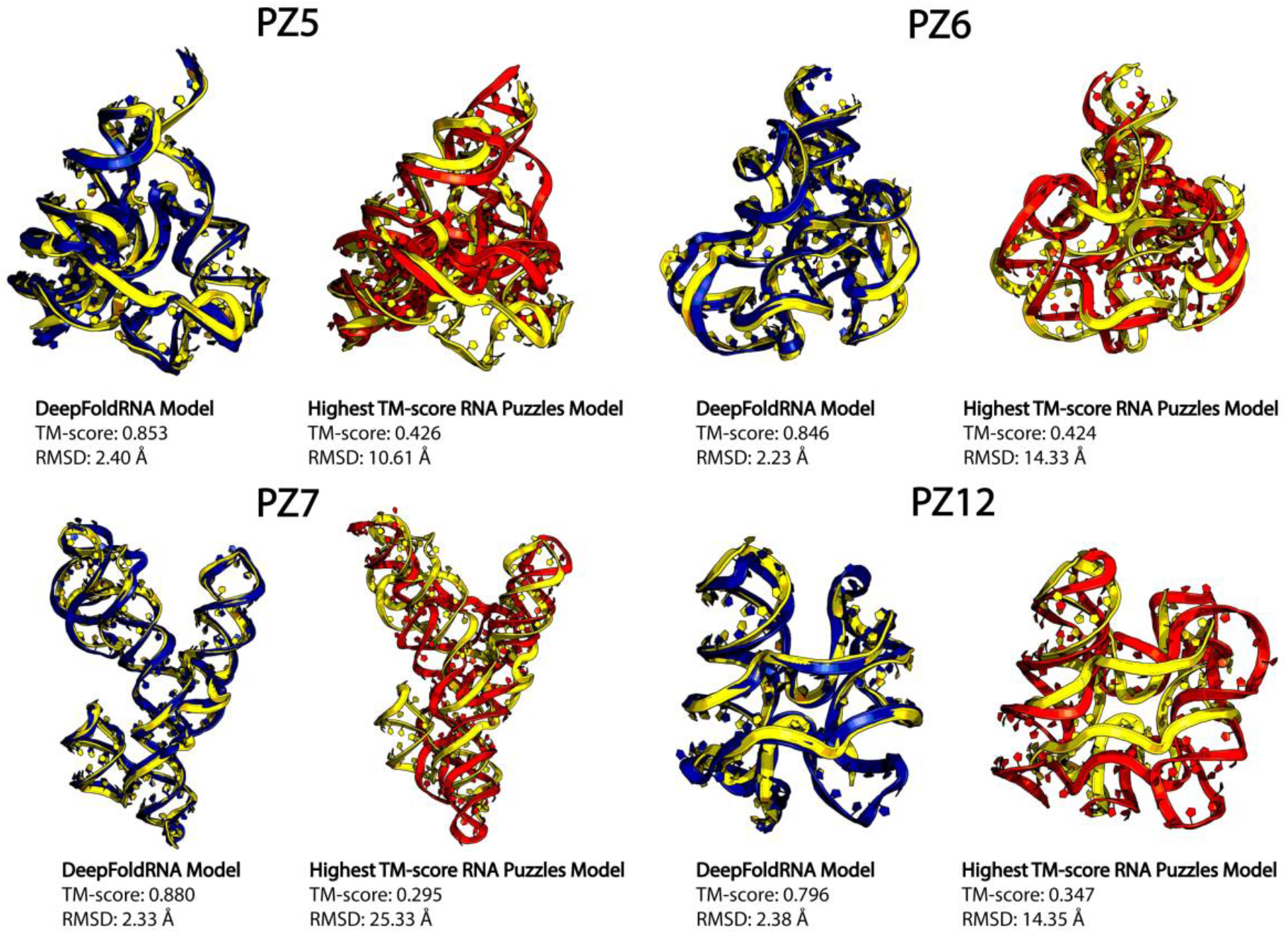
Case studies from difficult RNA-Puzzles targets, where the native structures are shown in yellow, and the structures by DeepFoldRNA and the best RNA-Puzzles models are shown in blue and red, respectively.

Second, **Puzzle 6** is a 168-nucleotide adenosylcobalamin riboswitch (PDB ID: 4gxy), which possesses a large ligand binding pocket that binds the adenosyl moiety to control gene expression^23^. The models submitted by the RNA-Puzzles community had a wide range of TM-scores (~0.142-0.424) and RMSDs (~38.02-11.89 Å), where the best model (TM-score=0.424) was produced with the assistance of experimental SHAPE data to help elucidate important secondary structure and contact information^13^. For DeepFoldRNA, a reliable MSA was collected with a high Neff of 517.9, which resulted in accurate predicted restraints with MAEs for the top 5*L* N-N distances and *Ω/λ* orientations of 0.94 Å and 0.22/0.17 radians, respectively. Moreover, the folding simulations produced a structure that closely matched the predicted restraints with an MAE of 0.73 Å between the top 5*L* predicted N-N distances and the model distances. Thus, the generated model possessed a near-native structure with a TM-score of 0.846 and RMSD of 2.23 Å. Importantly, the ligand binding site, which is essential to the RNA’s function, was accurately recapitulated without any explicit provisions or simulations that accounted for the ligand position.

Third, **Puzzle 7** is the Varkud satellite ribozyme (PDB ID: 4r4v), which is composed of 185 nucleotides and mediates rolling circle replication of a plasmid in the *Neurospora* mitochondrion^24^. The highest TM-score RNA-Puzzles model was constructed with the assistance of hydroxy radical footprinting experiments as well as mutate-and-map measurements used to determine contact information^6^. Nevertheless, the resulting model had a low TM-score of 0.295 and a high RMSD of 25.33 Å, where the model possessed incorrect helical orientations and an overly compact structure. For this target, DeepFoldRNA generated a poor MSA containing only 3 sequence homologs, all of which were nearly identical to the query sequence, resulting in an extremely low Neff of 0.07, making the prediction essentially a single sequence prediction problem. Nevertheless, the deep learning module produced accurate restraints with MAEs for the top 5*L* N-N distances and *Ω/λ* orientations of 0.77 Å and 0.16/0.14 radians, respectively. This resulted in a DeepFoldRNA model with a TM-score of 0.875 and RMSD of 2.40 Å, corresponding to a 196.6% higher TM-score than the best model submitted during RNA-Puzzles.

Last, **Puzzle 12** is a medium-size (108 nucleotides) *ydaO* riboswitch (PDB ID: 4qlm) with a novel structural topology that contains two binding pockets for cyclic-di-AMP^25^. It is involved in a number of important cellular functions, including sporulation, osmotic stress responses, and cell wall metabolism^25^. The best RNA-Puzzles model was produced with the assistance of fast-track experimental SHAPE data and multidimensional chemical mapping^6^ and had a TM-score of 0.347 and RMSD of 14.35 Å. Notably, the bubble region in the structure was unable to be correctly predicted by any of the submitted models and is partially unresolved in the crystal structure, likely due to its flexibility and lack of base pairing^6^. For DeepFoldRNA, the generated MSA was reliable with a Neff value of 135.5 and the resulting predicted restraints were accurate with MAEs of 0.70 Å and 0.15/0.20 radians for the top 5*L* N-N distances and *Ω/λ* orientations, respectively. Again, the folding simulations closely converged to the predicted restraints with an MAE of 0.72 Å between the predicted top 5*L* N-N distances and the model distances. These resulted in a high-quality model by DeepFoldRNA with an RMSD of 2.38 Å and a TM-score of 0.796 to the experimental structure, corresponding to a 129.4% improvement in the TM-score over the best model produced during the RNA-Puzzles challenge.

The results on these case studies demonstrate that DeepFoldRNA is able to produce accurate structural models for challenging RNAs that could not be folded by any traditional approach even with expert intervention and experimental restraints. It is practically encouraging that medium to higher resolution structures could be created for complex folds with few homologous RNA sequences, which has been one of the most challenging problems for deep learning-based protein structure modeling methods^15,17,26^. This is probably due to the fact that, compared to proteins whose structural patterns are often buried in deep evolutionary profiles, RNA structures are more explicitly encoded in the individual nucleic acid sequences (e.g., the tertiary structures are highly dependent on the Watson-Crick pairing of the RNA sequence), which can be readily captured by advanced deep learning models even with relatively shallow sequence profiles.

### DeepFoldRNA improves the speed and accuracy of RNA folding simulations for large RNAs

Monte Carlo sampling is a widely used approach in structural folding simulations and has been proven to be effective at identifying global free-energy minima for cases with frustrated knowledge-based energy landscapes^4,27,28^. However, these simulations typically require lengthy runtimes, which partially limits their application to large-scale modeling experiments. Given that the DeepFoldRNA energy landscape is significantly simplified by accurate and abundant spatial restraints, gradient-based L-BFGS sampling is sufficient to quickly fold RNA molecules and drastically reduce the simulation runtime (see Supplementary Video 1 for a real-time folding example).

As evidence, we plot in Figure 6A the simulation time required for DeepFoldRNA, SimRNA and FARFAR2 against the RNA length, where both SimRNA and FARFAR2 use Monte Carlo sampling. Overall, SimRNA required 379.3 minutes on average to fold the RNAs in the Rfam dataset, while DeepFoldRNA required 1.1 minutes, corresponding to a 345-fold reduction in the folding simulation time. The difference was even more significant when compared to FARFAR2, which required 4547.1 minutes for its folding simulations on average. Notably, DeepFoldRNA could fold the largest RNA in the dataset, which was composed of 237 nucleotides, within 7 minutes, while SimRNA and FARFAR2 required 1146 and 11615 minutes, respectively. Thus, DeepFoldRNA can be used to fold RNA molecules in seconds to minutes, significantly improving the speed at which RNAs can be modeled.

**Figure 6.**
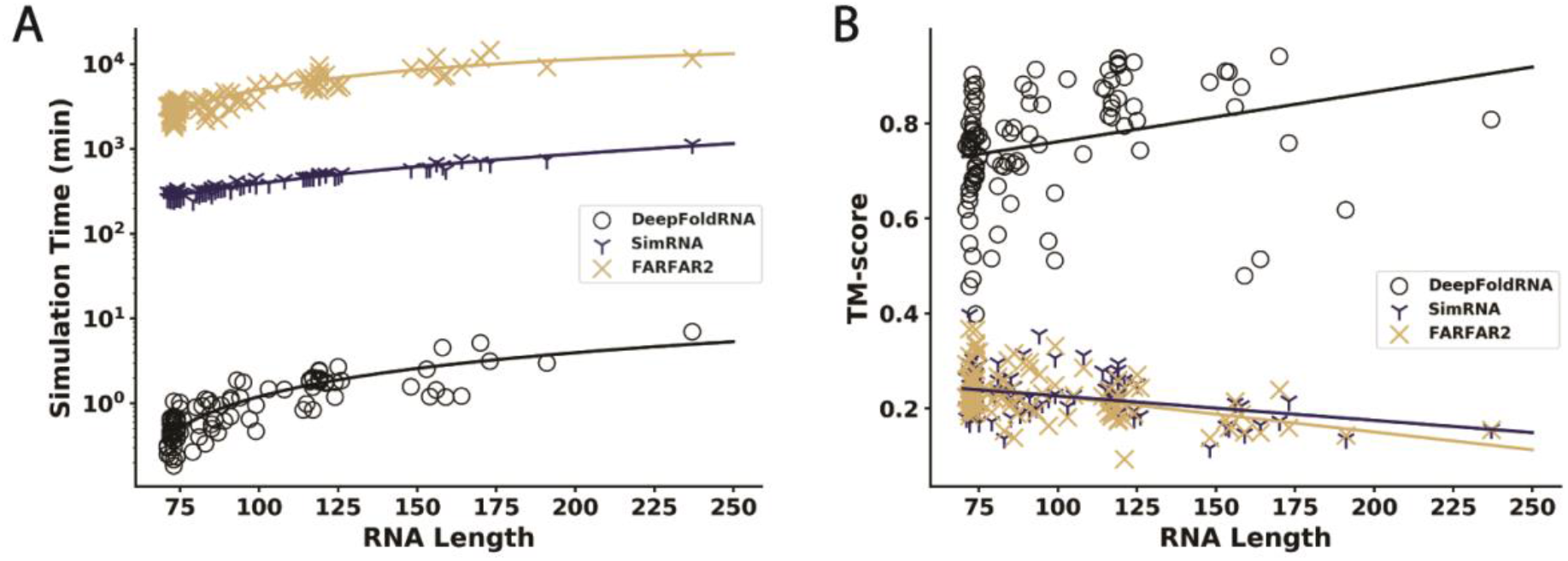
Dependence of the simulation runtime/modeling performance on the RNA length for DeepFoldRNA, SimRNA, and FARFAR2. A) Log-scale simulation runtime for DeepFoldRNA, SimRNA, and FARFAR2 in minutes plotted against the RNA length. B) Model TM-score versus RNA length for DeepFoldRNA, SimRNA, and FARFAR2. Lines are plotted to guide the eye.

Crucially, the modeling performance of DeepFoldRNA did not deteriorate as the sequence length of the RNA increased. In Figure 6B, we plot the TM-score values for the models generated by DeepFoldRNA, SimRNA, and FARFAR2 against the RNA sequence length. As expected, there is a negative correlation between the RNA length and model TM-score for both SimRNA and FARFAR2, as larger RNAs often have more complex folds that require sampling from wider-ranging conformational space, which is more difficult for Monte Carlo sampling to cover when guided by low-resolution energy force fields. For DeepFoldRNA, however, there is actually a slight positive correlation, where the method was able to generate on average more accurate spatial restraints and reliable folds for longer RNAs in the test set. These data demonstrate that the rapid simulations do not lead to unreliable results for larger and more complex folds, making DeepFoldRNA a robust method for generating accurate models independent of the fold complexity, which is critical for applications to large-scale RNA structure modeling.

## CONCLUSION

We developed a fully-automated method, DeepFoldRNA, to model RNA structures starting from sequence alone. The approach is built on deep self-attention neural networks to deduce high-accuracy spatial restraints from multiple RNA sequence alignments, followed by full-length 3D model construction through restraint-guided L-BFGS folding simulations.

The method was tested on two independent benchmark datasets. The first consisted of 105 non-redundant RNAs from 32 Rfam families with complex global folds. For these targets, DeepFoldRNA generated models with an average TM-score of 0.757 and RMSD of 2.68 Å, which was dramatically more accurate than the state-of-the-art methods, SimRNA^4^ and FARFAR2^5^, which are physical and knowledge-based Monte Carlo approaches that produced models with an average TM-score/RMSD of 0.228/19.37 Å and 0.228/21.07 Å, respectively. For the second benchmark dataset containing 17 challenging targets from the community-wide RNA-Puzzles experiment, DeepFoldRNA constructed higher quality models than the best models submitted from the community for 15 cases, where there was a large margin in the TM-score/RMSD difference for 10 cases, despite the fact that many models from the community were constructed with human expert intervention and experimental restraints^6,7,13^.

These improvements demonstrate the power and advantage of deep self-attention neural networks, which can learn more detailed structural information from evolutionary profiles than knowledge-based potentials derived from simple statistics of PDB structures. Nevertheless, the success of DeepFoldRNA modeling exhibited little correlation to the quality of the input MSAs, in part due to the effectiveness of self-attention networks, which are able to learn structural patterns embedded in single RNA sequences. Meanwhile, given the abundant, high-accuracy restraints produced by the deep learning modules, which can dramatically simplify the energy landscape, an additional advantage of DeepFoldRNA is its rapid model construction enabled by the gradient-based folding simulations. On average, DeepFoldRNA only required 1.1 minutes to fold the Rfam RNAs, which was 345 times faster than SimRNA and 4134 times faster than FARFAR2.

The high accuracy and speed of DeepFoldRNA, together with its fully-automated procedure, should help facilitate its usefulness and application to large-scale, atomic-level RNA structure modeling. Currently, only 2% of Rfam families have experimentally solved structures, where the application of DeepFoldRNA to model unknown Rfam families will provide critical information and insight into uncharacterized RNA structure space. Furthermore, the extension of the deep neural network models to RNA complexes and RNA-protein interactions will help elucidate the molecular and cellular functions of non-coding RNAs. Studies along these lines are in process.

## METHODS

DeepFoldRNA is a deep learning-based approach to full-length RNA structure modeling, which consists of three main steps: input feature generation, spatial restraint prediction, and L-BFGS folding simulations, as depicted in Figure 1.

### Input feature generation

DeepFoldRNA takes as input the nucleic acid sequence in FASTA format for the RNA of interest, from which all features used by the neural network are derived. The major input to the network is the MSA generated by rMSA^18^. Briefly, rMSA constructs an MSA for a query sequence by iteratively searching multiple nucleic acid sequence databases, including Rfam^3^, RNAcentral^2^, and the nt database^29^ using blastn^30^, nhmmer^31^, and cmsearch^31^ (Figure S4). From the MSA, nhmmer is used to generate a hidden Markov model (HMM), which serves as a succinct statistical representation of the detected homologous sequence profile. Additionally, PETfold^32^ is used to predict the secondary structure from the generated MSA, where the pairwise reliability scores are used as the input of the network. This allows for a convenient embedding of the secondary structure information, while simultaneously capturing the uncertainty in the PETfold predictions.

### Network Architecture

#### Overall architecture

The overall network architecture is depicted in Figure 1A. The generated features are first embedded into the network and passed through 48 MSA Transformer blocks, which use multiple self-attention layers to extract information encoded in the MSA to determine the spatial relationships between each pair of positions. Meanwhile, a pair representation is embedded into the network, where communication is encouraged between the MSA and pair representations using biased self-attention as well as updating the pair representation based on the processed MSA representation. Following this step, the sequence embedding is extracted from the MSA representation based on the position in the MSA embedding that corresponds to the original query sequence. The sequence embedding is then processed by 4 Sequence Transformer blocks, which use multiple self-attention layers that are biased by the pair representation to encourage consistency between the two. This process is repeated for 4 cycles, where the MSA and pair representations determined from the MSA Transformer layers of the previous cycle as well as the sequence embedding from the previous Sequence Transformer layers are added to those produced by the current iteration, allowing the network to gradually refine its predictions. Finally, the distance and orientation restraints are predicted from a linear projection of the final pair representation, while the backbone torsion angles are predicted from a linear projection of the sequence representation. A more detailed description of each component of the network is described below.

#### Input embedding

As shown in Figure 1A, the input features are used to generate two major representations: the MSA representation and the pairwise representation. The MSA embedding captures the evolutionary information contained in the MSA, while the pairwise representation captures the pairwise spatial relationships between each nucleic acid in the target sequence. To initialize the MSA representation, up to 128 sequences from the MSA are randomly sampled and encoded using one-hot-encoding. Then a linear layer with an output channel size of 32 is used to embed the one-hot-encoded MSA along with the relative positional encodings of each MSA column. The nhmmer HMM is also embedded using a linear layer with an output channel dimension of 32 and concatenated to the MSA embedding to produce the initial MSA representation. The pairwise representation is initialized from the paired sequence encodings as well as the predicted secondary structure using linear layers with an output dimension of 32. Here, the predicted secondary structure is not in dot-bracket format, but rather the *L*x*L* reliability scores output by PETfold, which allows for a convenient projection of the secondary structure information as well as inherently capturing the uncertainty in the predictions. A triangular self-attention procedure^15^ is applied to further refine the pair representation derived from the predicted secondary structure information. Triangular self-attention represents the pair embedding as a directed graph and performs multiple rounds of self-attention-based transformations by first updating the outgoing edges of the pair representation graph followed by the incoming edges. Then self-attention is performed around the starting nodes and around the ending nodes. Lastly, a transition block composed of two linear layers is used to project the pair embedding to an output dimension of 32*2 and back down to the original size of 32.

#### MSA Transformer network

The MSA Transformer network takes as input the MSA and pair embeddings, where the MSA representation contains information from homologous sequences in each row and the positional relationships in each column and the pair representation contains the pairwise distance relationships. The MSA embedding is first processed using multi-head, row-wise self-attention, which extracts positional information encoded by the different homologous sequences contained in the MSA. During the row-wise self-attention procedure, the rows of the MSA are mapped to a set of queries (*q*), keys (*k*), and values (*v*) using linear layers with an input dimension of 32 and an output dimension of 8×16, where 8 is the number of heads and 16 is the size of the hidden dimension.

The attention maps can be derived from the set of queries, keys, and values following the standard formulation: 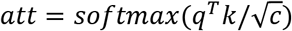, where *c* is the size of the hidden dimension. Bias from a linear projection of the pair representation is added to the resulting attention maps and the updated MSA row embeddings are determined by applying the attention maps to the values (*v*) along with a gate determined from a linear projection of the rows of the MSA representation followed by a sigmoid activation. After the row-wise self-attention layer, a similar procedure is repeated for the MSA columns using multi-head, column-wise self-attention. During this process, the columns of the MSA are mapped to queries, keys, and values using linear layers with an input dimension of 32 and an output dimension of 8×16, where 8 is the number of heads and 16 is the size of the hidden dimension. Next, the MSA columns are updated using the queries and keys to calculate the attention maps and applying these to the obtained values. Similar to the MSA row-wise self-attention, a gate is applied to the updated columns using a linear projection of the MSA columns followed by a sigmoid activation. The updated MSA rows and columns are further processed using an MSA transition block that passes the embedding through two linear layers, where the first layer projects the MSA embedding with a hidden dimension of 32 to a hidden dimension of 32×16 and the second layer projects the MSA embedding back to its original size of 32. Finally, the pairwise representation is updated by taking the outer product mean of the processed MSA representation and adding it to the pair representation. The pair representation is then processed using the same triangular self-attention scheme introduced above.

Overall, this process is repeated 48 times to gradually refine the MSA and pair embeddings, where the final output is the updated MSA and pair representations. If it is not the first cycle through the network, the MSA and pair representations from the previous pass of the network are then added to the representations from the current cycle.

#### Sequence Transformer network

Following the MSA Transformer layers, the position in the MSA embedding that corresponds to the original sequence is extracted and a linear layer is used to project its dimension from 32 to 64. Next, the sequence embedding is processed by two self-attention blocks. The first maps the input sequence to a set of queries, keys, and values from which the attention maps are derived. Bias from a linear projection of the pair representation is then added to the attention maps and the attention maps are applied to the values to update the sequence representation. A gate is also applied to the updated sequence representation by a linear projection of the sequence embedding followed by a sigmoid activation. The second attention block is similar to the first with the exception that it does not include bias from the pair representation. Finally, the updated sequence embedding is passed through 3 linear layers with an input and output channel dimension size of 64 to produce the final sequence representation. If it is not the first cycle through the network, the sequence representation from the previous pass through the network is added to the final sequence embedding from the current iteration.

#### Geometric restraint prediction

The predicted geometric restraints include the pairwise distance maps between the N1/N9 atoms, C4’ atoms, and backbone P atoms, as well as the inter-residue and backbone torsion angles (*ω, λ, η θ*) specified in Figure S1. The distances are divided into 40 bins, where the first and last bins indicate predicted distances <2 Å or >40 Å, respectively, while the middle 38 bins correspond to distances in the range of [2Å, 40Å] with an even bin width of 1Å. Similarly, the inter-residue torsion angles (*Ω, λ*) are divided into 25 bins with a width of 15°, where the first 24 bins correspond to the probability that the orientation angles fall in the range [-180°, 180°] and the last bin captures the probability that there is no interaction between a given pair of nucleotides. Non-interacting nucleotides are defined as those with N1/N9-N1/N9 distances >40 Å. The distance and orientation restraints are predicted from the final pairwise representation using linear layers with an input dimension of 32 and an output dimension of 40 for the distance restraints and 25 for the inter-residue orientation restraints, where a log softmax activation is applied to each output restraint. Lastly, the backbone pseudo-torsion angles (*η, θ*) are predicted from a linear projection of the sequence embedding with an input channel dimension of 64 and an output channel dimension of 24. Thus, the predicted pseudo-torsion angles are divided into 24 bins from [-180°, 180°] with a width of 15°, where a log softmax activation function is applied to the final predictions.

### Training Data and Procedure

DeepFoldRNA was trained on 2,986 RNA chains collected from the PDB, which were non-redundant (with a sequence identity <80%) to the 122 test RNAs used in this study. The labeled features from the PDB structures include the native C4’, N1/N9 and P distance maps, the inter-residue Ω and *λ* orientations, and the backbone *η* and *θ* pseudo-torsion angles. The training features from the experimental structures were discretized into binned values with the same sizes as the predicted features. The output of the network is the probability that each feature falls within one of the given bins, thus the network was trained using the softmax cross-entropy loss between the predicted and native distributions. In addition, a BERT-style loss^33^ was incorporated by randomly masking positions in the MSA and predicting the masked positions from a linear projection of the final MSA representation. The softmax cross-entropy loss was then calculated between the predicted MSA and the unmasked MSA.

Since the number of solved RNA structures is low, we also collected a non-redundant distillation set of 16,842 RNAs from the bp-RNA-1m database^34^ that were predicted to have regular secondary structures. Since the RNA in the distillation set do not have solved tertiary structures, we generated predicted labels for each RNA based on the network that was trained on the PDB sample. We then trained the network by sampling from the PDB dataset at a probability of 25% and from the distillation set at a probability of 75%. The loss function for the distillation set was identical to that used for the PDB set, where the softmax cross-entropy loss was calculated between the predicted features and the labels.

The network model was trained using Adam optimization with a learning rate of 0.001 for 159,000 epochs, where the distillation set was incorporated after epoch 99,000. The entire model was trained using a single Nividia V100 SMX2 GPU on the SDSC Expanse cluster^35^.

### DeepFoldRNA Energy Function

DeepFoldRNA uses L-BFGS simulations to quickly fold RNAs based on optimization of the following energy function:

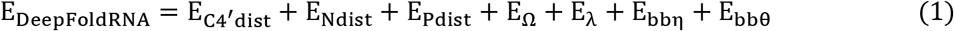

where E_C4’dist_, E_Ndist_, E_Pdist_, E_Ω_, E_φ_, E_bbη_, and E_bbθ_ are energy terms derived from the predicted C4’ – C4’ distances, N1/N9-N1/N9 distances, P-P distances, Ω orientations, λ orientations, backbone η torsions, and backbone θ torsions, respectively. The details of each energy term are further explained in Text S3. As L-BFGS optimization requires a continuously differentiable energy function, the energy terms are fit using cubic spline interpolation.

Overall, the DeepFoldRNA force field consists of 7 weighting parameters, which were determined on 184 RNAs from the training set with lengths between 60-480 nucleotides and pairwise sequence identities <50%. The weights were tuned iteratively, where the weight for each energy term was adjusted one at a time in the range [0, 10] using an increment of 0.25. The weighting parameter for each term that produced models with the highest average TM-score for the 184 RNAs were selected. Once the initial weight for each energy term was determined, this process was repeated 4 more times, varying each weight one-at-a-time using an increment of 0.1.

### L-BFGS Folding Simulations

In DeepFoldRNA, an RNA structure is represented by its backbone P and C4’ atoms as well as the N1/N9 atoms and two carbon atoms from the base (C2/C4 for pyrimidines and C2/C6 for purines) (Figure S1A). During the simulations, the bond lengths and bond angles are fixed at their ideal values, and the optimization is directly carried out on the backbone *η* and *θ* pseudo-torsion angles guided by the gradient of the energy function with respect to *η* and *θ*. L-BFGS optimization is used to find the backbone *η/θ* angles for each residue that minimize the energy function described in Eq. (1).

Here, L-BFGS is a gradient descent-based optimization method built on a limited memory variant of the Broyden-Fletcher-Goldfarb-Shanno (BFGS) algorithm, which attempts to identify the minimum of *E*_DeepFoldRNA_(*η, θ*), where *η/θ* are vectors of length *L* that represent the backbone pseudo-torsion angles at each position of the simulated structure. At each L-BFGS step *k*, the search direction *d_k_* is calculated by

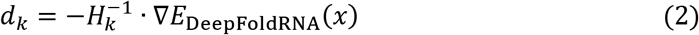

where 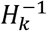 is an estimate for the inverse Hessian matrix and ∇*E*_DeepFoldRNA_(*x*) is the gradient of the DeepFoldRNA energy function with respect to the backbone pseudo-torsion angles, that is *x* = (*η, θ*). 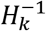 at step *k* = 0 is set to the identity matrix, *I*, and the value of 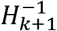 is obtained following the BFGS formulation:

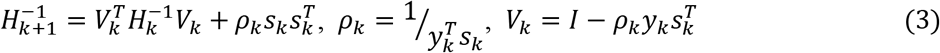

where *s_k_* = *x*_*k*+1_ – *x_k_* and *y_k_* = ∇*E*_DeeepFoldRNA_(*x*_*k*+1_) – ∇*E*_DeeepFoldRNA_(*x_k_*). Accordingly, the value of 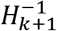 can be computed recursively by storing the previously calculated values of *s_k_* and *y_k_*. However, to preserve memory, L-BFGS only stores the last *m* values of *s_k_* and *y_k_*. Thus, *H*_*k*+1_ can be calculated as follows:

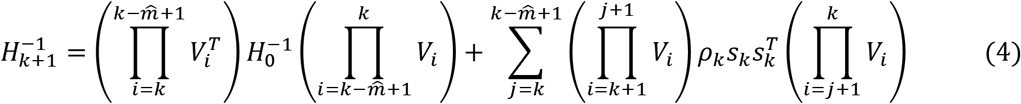

where 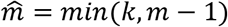 and *m* is set to 256 in DeepFoldRNA. Once the search direction *d_k_* is calculated, the *η/θ* angles are updated according to:

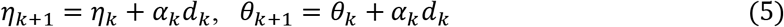

The value of *α_k_* is determined using the Armijo line search technique^36^ and dictates the amount to move along the given search direction. In DeepFoldRNA, a maximum of 10 L-BFGS iterations are performed with 2000 steps each, or until the simulations converge. We also use 3 rounds of noisy restarts, where the optimal backbone pseudo-torsion angles from the previous simulation are perturbed by a random value in the range [-10°, 10°] to avoid becoming trapped in local minima. The final model is the lowest energy decoy produced during the folding simulations.

## ACKNOWLEDGMENTS

This work used the Extreme Science and Engineering Discovery Environment (XSEDE^35^), which is supported by the National Science Foundation (ACI-1548562). This work was supported in part by the NIGMS (GM136422, S10OD026825), NIAID (AI134678), NIEHS (P30ES017885), NCI (U24CA210967), and the NSF (IIS1901191, DBI2030790, MTM2025426).

## AUTHOR CONTRIBUTIONS

Y.Z. conceived and designed the research; R.P. developed DeepFoldRNA, performed the experiments, analyzed the data, and constructed the webserver; R.P., G.S.O., and Y.Z. wrote the manuscript.

