## Supplemental Information for "*De Novo* RNA Tertiary Structure Prediction at Atomic Resolution Using Geometric Potentials from Deep Learning"

#### Supplementary Figures

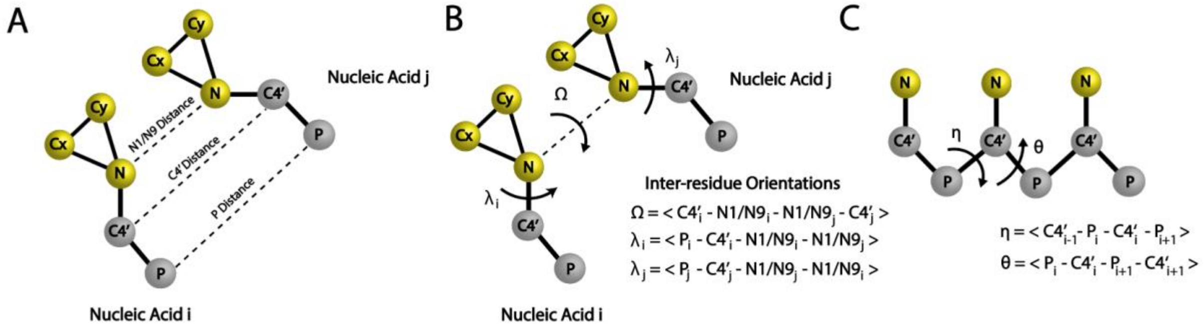


**Figure S1.** Definition of the geometric restraints predicted by DeepFoldRNA. including (A) inter-residue distances; (B) inter-residue torsion angles; and (C) backbone pseudo-torsion angles.

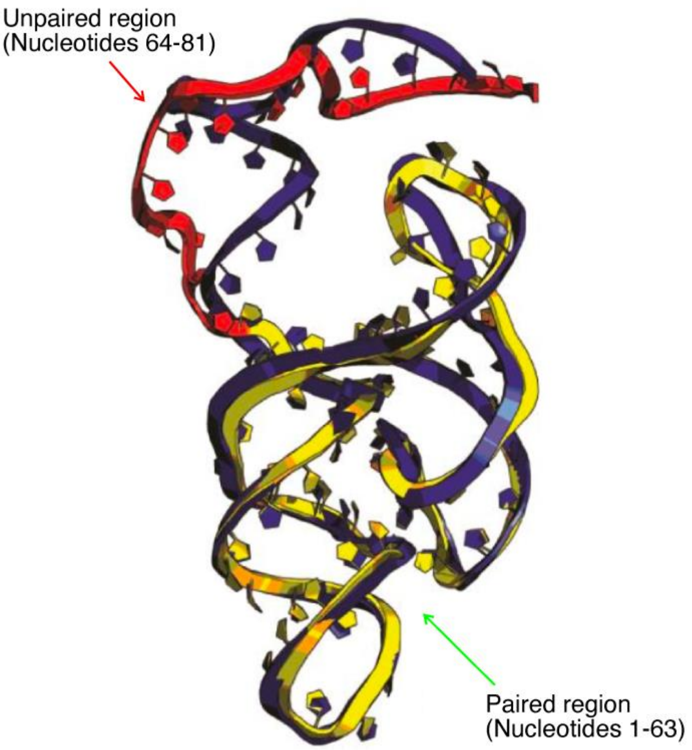


**Figure S2.** Case study from Rfam RNA RF02678, where the DeepFoldRNA predicted model (blue cartoon) is superimposed on the experimentally solved structure (PDB ID: 6jq5, chain A, nucleotides 1-81). The unpaired region in the experimental structure is shown in red and the paired region in yellow.

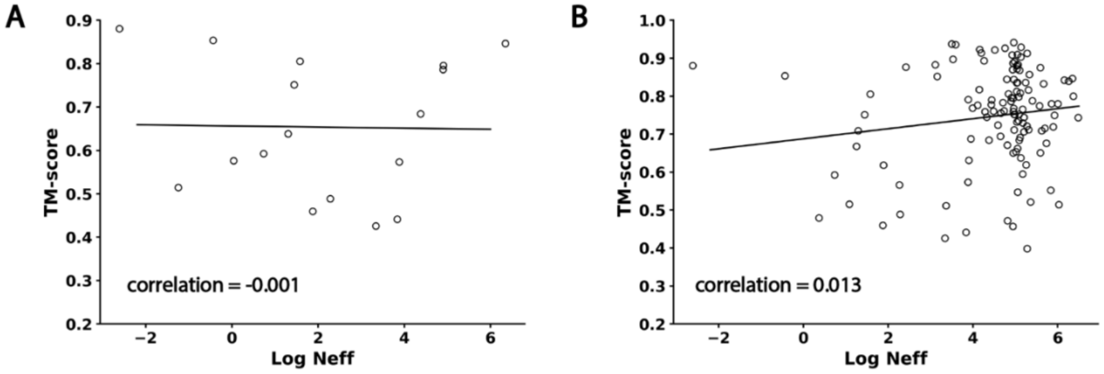


**Figure S3:** Model TM-score vs. the logarithm of the MSA Neff value for DeepFoldRNA on the RNA-Puzzles dataset (A) and the overall dataset (B), where the fitted models were obtained by linear regression.

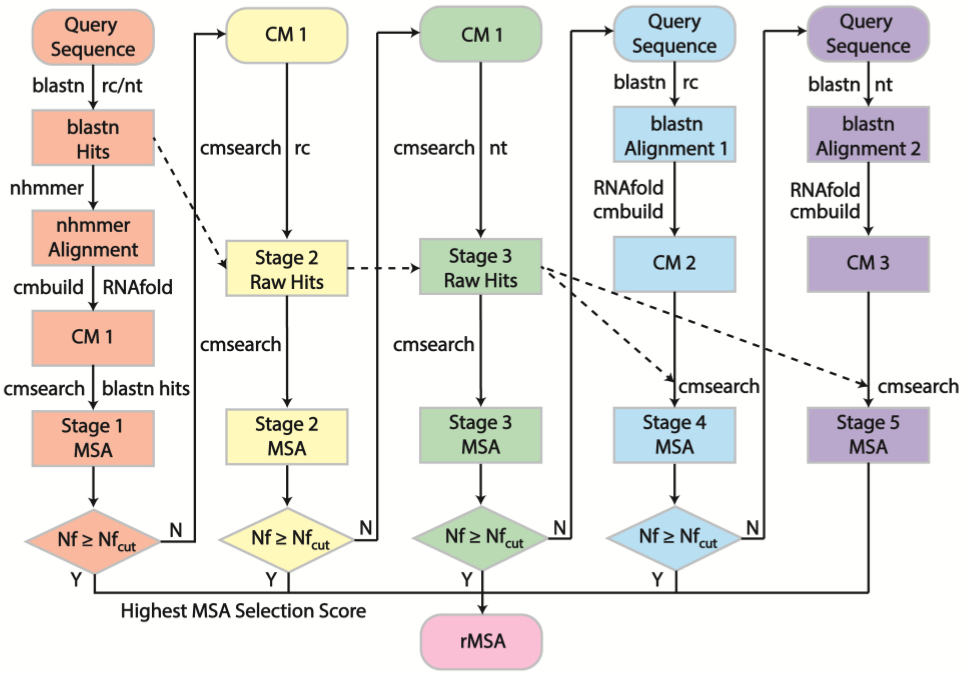


**Figure S4.** rMSA pipeline. rMSA generates 5 MSAs in total, where "CM" and "rc" stand for Covariance Model and the RNAcentral database, respectively, and $Nf_{\mathrm{cut}}$=128. The blastn searches are performed with "-max_target_seqs 50000 -strand plus" and "-max_target_seqs -strand both" options to search only the plus strand through RNAcentral and both strands through the nt database, respectively. This is due to the fact that RNAcentral is a transcriptomics database, while nt is a genomics database. Similarly, cmsearch is performed using the "--toponly --incE 10.0" option for the plus strand in RNAcentral and "--incE 10.0" for both strands in nt. The nhmmer search is performed using "--watson" to only consider alignments with directions that are consistent with the blastn alignments.

### Supplementary Texts

**Text S1: SimRNA Procedure.**
SimRNA was run using the following command:

```
<SimRNA_Directory>/SimRNA -s seq.fasta -c config.dat -S SecondaryStructure.txt
```

The default configuration file was used which runs 16,000,000 folding iterations, where the contents of the file are below:

```
NUMBER_OF_ITERATIONS 16000000
TRA_WRITE_IN_EVERY_N_ITERATIONS 16000

INIT_TEMP 1.35
FINAL_TEMP 0.90

BONDS_WEIGHT 1.0
ANGLES_WEIGHT 1.0
TORS_ANGLES_WEIGHT 0.0
ETA_THETA_WEIGHT 0.40
```

The final model was selected from the lowest energy decoy generated from each simulation.

**Text S2: FARFAR2 Procedure.**
FARFAR2 was run using the following command:

```
<Rosetta_bin>/rna_denovo.static.linuxgccrelease -fasta seq.fasta -native native.pdb -out:file:silent out.txt -nstruct 100 -minimize_rna true -fragment_homology_rmsd 1.2 -secstruct <Secondary Structure>
```

The default number of cycles were run for each simulation (10,000) and the final model was selected following clustering of the 100 generated structures using the default cluster radius of 3 Å and selecting the first cluster as the representative model.

**Text S3: DeepFoldRNA energy function.**
The energy function used to guide the DeepFoldRNA simulations is a linear combination of 7 energy terms:

$$\mathrm{E_{DeepFoldRNA}} = \mathrm{E_{C4'dist}} + \mathrm{E_{Ndist}} + \mathrm{E_{Pdist}} + \mathrm{E_{\Omega}} + \mathrm{E_{\lambda}} + \mathrm{E_{bb\eta}} + \mathrm{E_{bb\theta}} \tag{1}$$

where $\mathrm{E_{C4'dist}}$, $\mathrm{E_{Ndist}}$, $\mathrm{E_{Pdist}}$, $\mathrm{E_{\Omega}}$, $\mathrm{E_{\lambda}}$, $\mathrm{E_{bb\eta}}$, and $\mathrm{E_{bb\theta}}$ are energy terms derived from the predicted C4'–C4' distances, N1/N9-N1/N9 distances, P-P distances, Ω orientations, $\lambda$ orientations, backbone η torsions, and backbone θ torsions, respectively. All of the energy terms are based on pairwise interactions between residues *i* and *j* in an RNA molecule, with the exception of $\mathrm{E_{bb\eta}}$ and $\mathrm{E_{bb\theta}}$, which are single-body potentials. Thus, the cumulative terms are derived from the summation over all residue pairs i and j as follows:

$$E_{\mathrm{C4'dist}} = \sum_{i,j} E_{d_{ij}}(i,j) \tag{2}$$

$$E_{\mathrm{Ndist}} = \sum_{i,j} E_{d_{ij}}(i,j) \tag{3}$$

$$E_{\mathrm{Pdist}} = \sum_{i,j} E_{d_{ij}}(i,j) \tag{4}$$

$$E_{\Omega} = \sum_{i,j} E_{\Omega_{ij}}(i,j) \tag{5}$$

$$E_{\lambda} = \sum_{i,j} E_{\lambda_{ij}}(i,j) + \sum_{j,i} E_{\lambda_{ji}}(j,i) \tag{6}$$

$$E_{\mathrm{bb\eta}} = \sum_{i} E_{\mathrm{bb\eta_i}}(i) \tag{7}$$

$$E_{\mathrm{bb\theta}} = \sum_{i} E_{\mathrm{bb\theta_i}}(i) \tag{8}$$

Note, the inter-residue $\lambda$ orientation is not symmetric, thus it must be summed over residues pairs ($i$, $j$) as well as the opposite direction ($j$, $i$).

The detailed description of each energy term is defined as:

$$E_{d_{ij}}(i,j) = \begin{cases} -\log\left(\dfrac{P\left(d_{ij}\right) + \epsilon}{P(d_{cut}) + \epsilon}\right), & d_{ij} < d_{cut} \\ 0, & d_{ij} \geq d_{cut} \end{cases} \tag{9}$$

where $d_{ij}$ is the distance between two C4' atoms for the C4' distance restraints, two N1/N9 atoms for the N1/N9 distance restraints, or two P atoms for the P distance restraints from residues $i$ and $j$, $P\left(d_{ij}\right)$ is the predicted probability by DeepFoldRNA associated with the distance $d_{ij}$, and $P(d_{cut})$ is the probability for the final distance bin which corresponds to a distance between 39-40 Å. The pseudo count $\epsilon = 1E-4$ is used to avoid issues when $P(d_{cut})$ is small. Cubic spline interpolation is used to interpolate between the energy at the different distance bins in order to make the potential differentiable for L-BFGS optimization.

$$E_{\Omega_{ij}}(i,j) = \{-\log\left(P\left(\Omega_{ij}\right) + \epsilon\right) \tag{10}$$

$$E_{\lambda_{ij}}(i,j) = \{-\log\left(P\left(\lambda_{ij}\right) + \epsilon\right) \tag{11}$$

$$E_{\lambda_{ji}}(j,i) = \{-\log\left(P\left(\lambda_{ji}\right) + \epsilon\right) \tag{12}$$

where $\Omega_{ij}$ and $\lambda_{ij}$ are the inter-residue orientations predicted by DeepFoldRNA between residues $i$ and $j$ defined in Figure S1. Furthermore, given that $\lambda$ is not symmetric for a residue pair, $\lambda_{ji}$ is the inter-residue orientation between residues $j$ and $i$. The pseudo count $\epsilon = 1E-4$ is used to avoid issues when the predicted probability is small. Cubic spline interpolation is used to interpolate between the energy at the different orientation bins in order to make the potential differentiable for L-BFGS optimization.

$$E_{bb\eta}(i) = \{-\log(P(\mathrm{bb\eta_i}) + \epsilon) \ \ and \ E_{\mathrm{bb\theta}}(i) = \{-\log(P(\mathrm{bb\theta_i}) + \epsilon) \tag{13}$$

where $\mathrm{bb}\eta_i$ and $\mathrm{bb}\theta_i$ are the backbone pseudo-torsion angles predicted by DeepFoldRNA for residue $i$. The pseudo count $\epsilon = 1E - 4$ is used to avoid issues when the predicted probability is small. Cubic spline interpolation is used to interpolate between the energy at the different torsion bins in order to make the potential differentiable for L-BFGS optimization.
